# Task demands and visual context naturalness modulate gravitational expectation during ocular tracking of temporarily occluded ballistic trajectories

**DOI:** 10.1101/2025.07.03.662924

**Authors:** Sergio Delle Monache, Pietro Nastasi, Gianluca Paolocci, Francesco Scalici, Riccardo Ingrosso, Francesco Lacquaniti, Iole Indovina, Gianfranco Bosco

## Abstract

During ocular tracking of a target, contextual information provides useful cues to account for motion features the visual system is poorly sensitive to, like its acceleration. For example, the brain can rely on an internal model of gravity to intercept and track accelerated targets under gravity, and cues about the visual environment naturalness can facilitate its recruitment. Notably, in real life, these two tasks are often associated. Experimental evidence suggests that predictive information may be shared between the two tasks, albeit their concurrent execution can, in turn, affect the oculomotor performance. To gain further insight on how task demands and the naturalness of the visual context affect the weighting of sensory feedback and predictive feedforward information in ocular tracking, subjects tracked targets either congruent with gravity or perturbed during the trajectory with altered gravity. Targets were projected on either pictorial or neutral background, and, to enforce prediction, were occluded for variable intervals 550 ms after the perturbation. Sixty-nine participants were divided into two groups: one group performed only ocular tracking, the other also intercepted the targets. In general, subjects’ ocular tracking performance depended on the target acceleration level and the visual context, with effects of gravitational expectations being more evident with the pictorial naturalistic environment. This result reinforced the idea that gravity a priori information can be weighted based on the naturalness of the visual context. Another striking finding was that participants who only tracked the target showed systematic changes of the oculomotor performance with the acceleration level right after the perturbation, compatible with a strong reliance on gravitational expectation; instead, participants that also intercepted the target showed such systematic changes primarily after the target’s occlusion, perhaps weighting more sensory information until then. Thus, concurrence of ocular tracking and manual interception seemingly influenced the temporal recruitment of internalized gravity information.

## INTRODUCTION

Visual information is crucial to interact efficiently with the environment, particularly when surrounding objects are moving. In this case, besides the object’s location, also information about its kinematics and the potential physical interaction with other objects becomes essential. Acquisition of high-definition visual information about the moving target is afforded by eye movements that track its motion, bringing and maintaining its image on the fovea (1–3). In particular, tracking of a target moving on the fronto-parallel plane can be achieved by combining two types of eye movements: a) saccadic movements, that rapidly shift the gaze to bring the object’s image from the peripheral retina to the fovea; b) smooth pursuit eye movements, which, by moving the eyes at the same velocity as the visual object, stabilize the retinal projection of the moving object on the fovea (4). The time required for processing visual information and executing these eye movements, however, introduces potentially significant delays, which must be accounted for to ensure accurate tracking (5). In this respect, predictive mechanisms that anticipate the target future behavior can play a decisive role to overcome the delays inherent to the sensorimotor loops (6,7). These predictions become more predominant when target information is temporarily missing –e.g., because of a temporary visual occlusion or in case of large gaze shifts– as they encompass temporal windows exceeding greatly the mere duration of sensorimotor delays (8–11).

Predictive estimates of the target motion can be based on signals about the target kinematics immediately preceding the occlusion (12), integrated with cognitive factors and a priori information derived from past experiences (13–15). A priori information may include also factors related to the overall naturalness of the visual environment and long-term memory of the dynamic interactions between the target and the surrounding environment, such as bounces against surfaces or deflections (8,16–18). In this respect, gravity represents a major physical invariant of the Earth environment, imposing ∼9.81 m s^-2^ downward acceleration at sea level to free-moving targets, and contributing to the perceived naturalness of the environment (19). Over the last couple of decades, several studies have gathered evidence that an internal model of gravity effects on the objects’ motion may play a role during manual interceptive actions to predictively estimate the time to contact with objects under the effect of gravity (20– 24. For a review, see 25), or during visual search tasks (26) and during the processing of vestibular information (27–29). This internal model may account for the accurate temporal and spatial estimates driving the interception of free-falling accelerating targets, in spite of the relatively poor sensitivity of the visual system to accelerated motion (30–33). The putative neural correlates of the gravity internal model have been identified within the vestibular network, a complex of multisensory brain areas mostly located around the Sylvian sulcus (34–36). More recent experimental evidence has extended the notion of the internal model of gravity to the oculomotor control (8,16,18,37,38). Delle Monache et al. (2015) (39) hypothesized that a priori information about gravity effects on the object motion may be shared by both interceptive and oculomotor control, since ocular tracking of trajectories perturbed with effects of altered gravity reflected expectation of gravity effects in a similar fashion to what observed for interceptive movements of the same target trajectories. Alike interceptive movements, the relative weight of a priori and sensory information driving eye movements was shown to depend strongly on the visual context. In a later study, Delle Monache et al. (2019) (37) manipulated the visual context on which moving targets were presented and reported that participants relied more strongly on sensory information when tracking targets against a uniform background, while expectations of gravity effects were more evident when the same target trajectories were tracked in a quasi-realistic pictorial background. However, during periods of transient disappearance of the visual targets, a priori expectations of gravity effects were evident with both visual contexts.

In addition to the subjects’ expectations on the motion of the visual object, another factor that can influence the eye-tracking behavior is the simultaneous execution of a contingent task. Renowned experiments by Yarbus have shown that, during the mere exploration of a scene, eye movement trajectories relate to the aim of that exploration (40). Moreover, demanding cognitive tasks influence oculomotor performance, with a clear negative correlation between the complexity of the task and the eye movements accuracy (41–45). For example, saccades and smooth pursuit movements produced during forced eye-tracking are altered when cognitive tasks are performed simultaneously, with a general increase in saccadic latencies, and decrease in saccadic accuracy and smooth pursuit gains for both reflex and voluntary eye movements (44). Neuroimaging studies have suggested that this behavior may be explained by the partial overlap of the cortical regions involved in the cognitive and oculomotor tasks (46,47).

A behavior often associated with the ocular tracking of a moving object is the manual interception. Indeed, during interceptive actions, eye and hand movements are generally coupled (48) and this coupling does provide an advantage to the interceptive performance, with a clear increase in the interception accuracy (49). Eye motion generally tends to lead hand movements (39,48,50), preceding the hand motion onset even during a go/no-go interception task (51). Corollary signals related to oculomotor predictions when the target motion is predictable appear to contribute to the planning of the interceptive action, accounting, perhaps, for the improvement of the interceptive performance, even independently from the eye-tracking performance itself (52–55). Moreover, as mentioned earlier, control centers for oculomotor and manual interception may share predictive information related to the effects of gravity on the object motion, which can be engaged even by pictorial representations of a natural environment (37,39). Specifically, in one of these earlier studies, Delle Monache and colleagues (2015) (39) examined the eye movement patterns that participants produced while intercepting visual targets moving with laws of motion either congruent or not with natural gravity and that could be occluded or not for variable time intervals. They reported that eye movement patterns depended on targets’ laws of motion and visibility, thereby suggesting predictive mechanisms. Moreover, with occluded targets, better interceptive performance was associated to greater eye tracking accuracy, supporting the idea that precise ocular tracking could provide better predictions of the target motion for the interceptive response. In this study, however, participants could move freely their eyes around the visual scene, thereby producing idiosyncratic patterns which did not allow to evaluate systematically how the concurrent execution of the manual interceptive task influenced the ocular tracking performance.

In the present study, we took a different approach to investigate more in depth the degree to which ocular tracking performance could be influenced by the concurrent execution of a manual interception, also with respect to the use of shared a priori information about the effects of natural gravity on the object motion. For this purpose, two groups of subjects tracked visual targets, projected on either pictorial or neutral background, which moved either congruently or not with the effects of gravity and, to enforce predictive mechanisms, were occluded for variable time intervals.

One group of participants was required to perform only continuous ocular tracking of the visual target, while the other group had to perform concurrently continuous ocular tracking and manual interception of the visual target. Firstly, our results provided further support to the notion that both eye tracking and interceptive movements can reflect expectations of gravity effects, depending on the visual context and on the visibility of the target. Secondly, a remarkable novel finding emerged in that the concurrence of eye tracking and manual interception influenced the temporal course with which gravity internalized information was engaged.

## METHODS

Sixty-nine volunteers signed informed written consent to participate to the experimental procedures. A priori power analysis yielded a required sample size of 68 participants to detect an effect size f of 0.5 with α = 0.05 and power of 0.9. Participants’ vision was either normal or corrected-to-normal. Subjects were randomly divided into two groups: the first group (A) comprised 37 subjects (20 women, 17 men; mean age 23.88 years ± 2.87 SD); the second group (B) comprised 32 subjects (18 women, 14 men; mean age 24.22 years ± 3.52 SD). Volunteers belonging to group B –who were required to perform a manual interception task, in addition to the oculomotor task– were tested for handedness by the abbreviated version of the Edinburgh Handedness Inventory (56). The large majority (n=25) were right-handed, five were ambidextrous, the remaining two, albeit left-handed, preferred using the right hand to control the computer mouse (25, 5 and 2 respectively). Group A participants were not tested for handedness because they were required to perform only the oculomotor task.

We opted for a between-subject design, rather than a within-subject design, to study the potential effects of the concurrent execution of the ocular tracking and the manual interception tasks on the oculomotor performance, based on the following considerations: a between-subjects design could reduce the risk of contamination across conditions and help preserving the independence of task contexts, which is particularly important when tasks differ substantially in structure or content (57); conversely, a within-subjects design would have increased the duration of each experiment to over 3 hours, raising the issue of controlling for fatigue, attention loss, and carryover effects. Thus, overall, this design provided a balanced trade-off between the statistical constraints and the practical feasibility. Experimental procedures were performed in agreement with the Declaration of Helsinki and approved by the ethics committee of the University of Rome “Tor Vergata” (protocol n. 140.12).

### Visual scenes

Subjects seated 60 cm away from a 22-in. LCD screen (ViewSonic VX2268WM), with their head stabilized by a chin rest. We presented two different visual scenarios, designed by means of the software package Presentation (version 14.9; Neurobehavioral Systems, Berkeley, CA). Scenarios were displayed with a refresh rate of 100 Hz and a spatial resolution of 1,680 * 1,050 pixels.

One scenario, the Pictorial Scenario, represented a play from the baseball game and contained quasi-realistic graphic elements and depth cues. A visual target started its motion from a baseball hitter located at the bottom left corner of the baseball field (Figure 1A). During its quasi-parabolic trajectory (see below), the target was transiently occluded by a smoke-like cloud (occluder) positioned, from trial to trial, in different locations on the upper right area of the screen. For the manual interception task (see below), the position of an outfielder, initially located at the lower central portion of the scene, could be controlled by displacing the computer mouse along the horizontal axis. A semitransparent circle (22 pixels diameter, 0.62° visual angle) surrounded the outfielder’s right hand to delimit the valid interception area.

**Figure 1.**
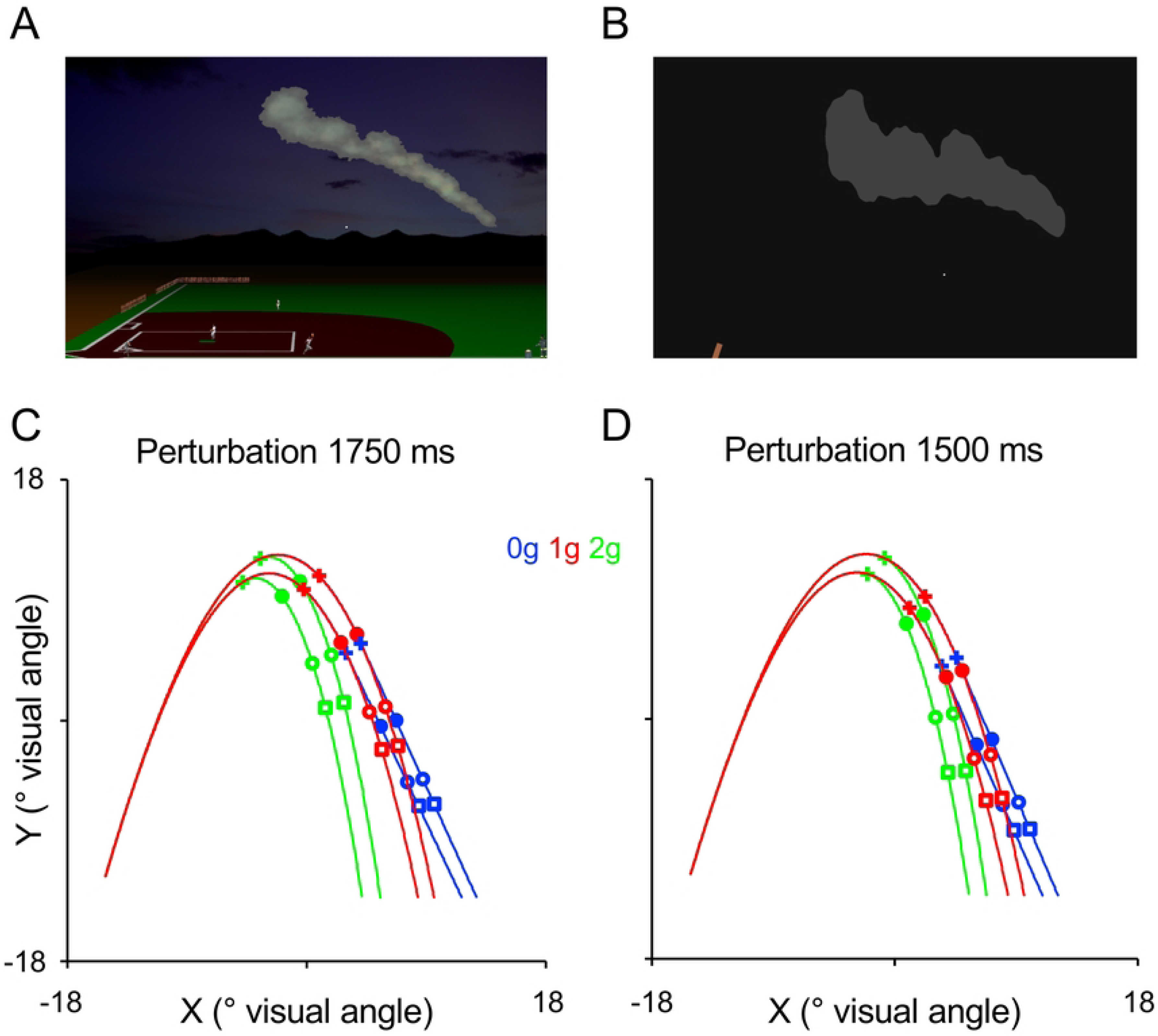
Visual scenes for the Oculomotor Task. A) In the Pictorial Scenario, the scene represented the fly-ball play of a baseball game. A white ball moved from the bottom left corner of the screen and followed a quasi-parabolic trajectory landing in the right portion of the screen. During its trajectory, the ball was transiently occluded by a smoke-alike cloud positioned in the upper right portion of the scene. Pictorial elements gave participants depth and relative sizes cues. B) In the Neutral Scenario, the white target moved on a dark-grey background with no graphic elements except for an orange rectangle on the lower left portion of the screen and an irregularly shaped light-gray geometric element on the upper right area of the scene. Visual scenes for the Oculo-manual Task were identical but with a white semitransparent 22 pixels circle on the glove of the outfielder in the Pictorial Scenario. The same graphic element was added also to the Neutral Scenario and in both cases represented the interception area of the interceptive task. C-D) Target trajectories. The ascending portion of the trajectory was modeled accounting for the effects of Earth gravity (scaled to the graphic elements of the Pictorial Scenario, see A). The descending portion of the trajectory could either retain the effects of gravity (1g, red) or be perturbed to simulate a micro-gravity (0g, blue) or a hyper-gravity (2g, green) condition. The + symbol indicates the point of the trajectory at which the kinematic perturbation was applied to produce 0g and 2g trajectories, while for unperturbed 1g trajectories it represents the temporal marker for determining the onset of the target occlusion (see the main text for details). 550 ms after the temporal marker of the perturbation, targets were temporarily occluded (filled circles) reappearing 450 ms (open circles) or 650 ms (open squares) later. C) trajectories perturbed 1750 ms before landing. D) trajectories perturbed 1500 ms before landing.

The other scenario, the Neutral Scenario, consisted of a dark-grey background (RGB colormap: 17, 17, 17) containing several geometrical elements. An orange rectangle represented the starting position of the target motion and a light gray irregularly-shaped geometric element occluded the target motion like the cloud of the Pictorial Scenario, to which it was matched for mean luminance (RGB colormap: 64, 64, 64; Figure 1B). In this scenario, the moving cursor for intercepting the visual target was a white circle (RGB colormap: 255, 255, 255) identical in size and starting position to the circle surrounding the outfielder’s hand in the Pictorial Scenario.

### Trajectories

Visual targets (white circles with a diameter of 7 pixels, 0.20° visual angle, RGB colormap: 255, 255, 255) followed ballistic trajectories confined to the frontal plane. Targets’ kinematics were in accordance with motion equations described by Brancazio (1985) (58) for a baseball fly ball affected by gravity and air drag (see also 21). Briefly, targets’ initial velocities could assume two possible values, instrumental to increase the variability in the experimental conditions (Figure 1C-D). The ascending portion of the trajectory accounted for the effects of the gravitational acceleration (g = 9.81 ms^-2^) scaled to the graphic elements of the Pictorial Scenario, whereas the descending portion could be either perturbed or not with altered gravity effects. Perturbations occurred at temporal markers of 1750 ms or 1500 ms before target landing: after these temporal markers, in 1/3 of the trials, targets retained gravity effects (1g conditions), while in the remaining 2/3 they could either assume constant velocity motion (0g conditions, 1/3 of the total number of trials) or increase instantly their constant acceleration to 2g (2g conditions, 1/3 of the total number of trials). Then, 550 ms after the perturbation marker, the target disappeared behind the occluder (the smoke-like cloud or the light-gray geometric element in the Pictorial and in the Neutral Scenario, respectively) to reappear either 450 ms or 650 ms later. Depending on the combination of perturbation and occlusion intervals, the target became visible again before landing for a time interval comprised between 300 and 750 ms. Alike the manipulations of the target initial velocity, the two perturbation intervals and the two occlusion intervals were introduced merely to increase the numerosity of the experimental conditions (i.e. increase the number of different trajectory types) and prevent that subjects’ oculomotor performance could reflect categorization of a reduced number of target trajectories. Moreover, in previous studies using similar experimental settings, we found that these manipulations could affect only marginally the oculomotor performance (37,39). Thus, overall, 24 experimental conditions were defined by combining 3 laws of motion (0g, 1g, 2g), 2 perturbation intervals (1750 ms, 1500 ms), 2 occlusion durations (450 ms, 650 ms) and 2 initial velocities. For each scenario, we presented 8 repetitions of each experimental condition (192 trials, overall), distributed pseudo-randomly. Trials with the two different scenarios were performed in separate blocks, with a 10-minute resting break between them. The order of the two scenarios was counterbalanced across subjects (Pictorial first and Neutral first).

### Behavioral tasks

As mentioned above, subjects were assigned randomly to two groups (A and B), performing different behavioral tasks. Subjects belonging to group A were required only to track continuously with their eyes the visual target throughout its trajectory, including the visually occluded portion (Oculomotor Task). Subjects belonging to group B performed concurrently ocular tracking and manual interception of the visual target (Oculo-Manual Task). In essence, subjects tracked the visual target continuously as group A subjects, while, at the same time, displacing with a gaming mouse (Razer Copperhead, San Francisco, CA) a cursor on the screen (i.e., the outfielder’s hand surrounded by the white circle in the Pictorial Scenario or the white circle in the Neutral Scenario) from its initial location towards the target landing location in the bottom right corner of the scene to intercept it. Subjects indicated the actual interception of the visual target with the white semitransparent circle by pressing a mouse button. As continuous ocular tracking of the target was imposed for both the Oculomotor and the Oculo-Manual task, we were able to compare directly the oculomotor performance between the two subject groups by using the same oculomotor indexes. It should be also noted that, although the manual interception task was identical to that described by previous studies of our group (21,39), the oculomotor requirements of the participants were profoundly different, because, unlike the current experiments, subjects were not constrained to tracking continuously the visual target, but they could move their eyes freely throughout the visual scene following patterns instrumental to the manual interception of the visual target.

### Data acquisition

Subjects’ binocular eye movements were recorded with an EyeLink 1000 tracker system at a sampling frequency of 500 Hz (SR Research, Ontario, Canada). The eye tracker was calibrated at the beginning of each experimental session and every 32 trials with a 9-point calibration grid, while drift corrections were applied every 8 trials. The mouse cursor position during the Oculo-manual Task was sampled at 200 Hz while button presses were sampled at 1 kHz.

### Eye movement analysis

Eye tracker signals from both eyes were analyzed preliminarily by means of a custom-made MATLAB (2022a; Mathworks, USA) script to detect blinking artifacts or missing data in the eye position traces. With this preliminary screening, we identified 299 (1.1%) out of 26,496 trials (the total number of trials in the experiment series resulting from 384 trials for each experiment multiplied by 69 subjects) that had to be discarded because signals from both eyes were not usable or missing. For 6,555/26,496 trials (24.7%), only signals from one eye were not usable or missing, but we retained data from the unaffected eye. For the remaining 19,642/26,496 trials with valid binocular eye traces, we computed the cyclopean eye trajectory by averaging the eye traces from the two eyes. Thus, overall, data from 26,197/26,496 trials (98.9%), obtained either from the spared trace of one eye or by averaging the signals from both eyes, were used for further analyses. Notably, the fraction of valid trials identified after the preliminary analysis was equal between the two subject groups (14,052/14,208, 98,9% for Group A and 12,145/12,288, 98,9% for Group B). Valid eye-tracker signals were numerically differentiated and filtered with a zero-lag 2nd-order low-pass Butterworth filter (cutoff frequency of 40 Hz, see 37,39,59). We evaluated subjects’ eye-tracking behavior during the two temporal windows illustrated in Figure 2: 1) the *perturbation* window (highlighted in *green*), beginning 100 ms after the perturbation temporal marker and ending at the visual occlusion onset 450 ms later; 2) the *occlusion* window (highlighted in *orange*), starting 100 ms after the target occlusion and ending 350 ms later (therefore including the entire occluded portion of the trajectory for the two occlusion durations). We did not consider the first 100 ms of data after the perturbation and occlusion events to take into account the visuomotor delays occurring between the incoming visual information and the oculomotor response (37,60).

**Figure 2.**
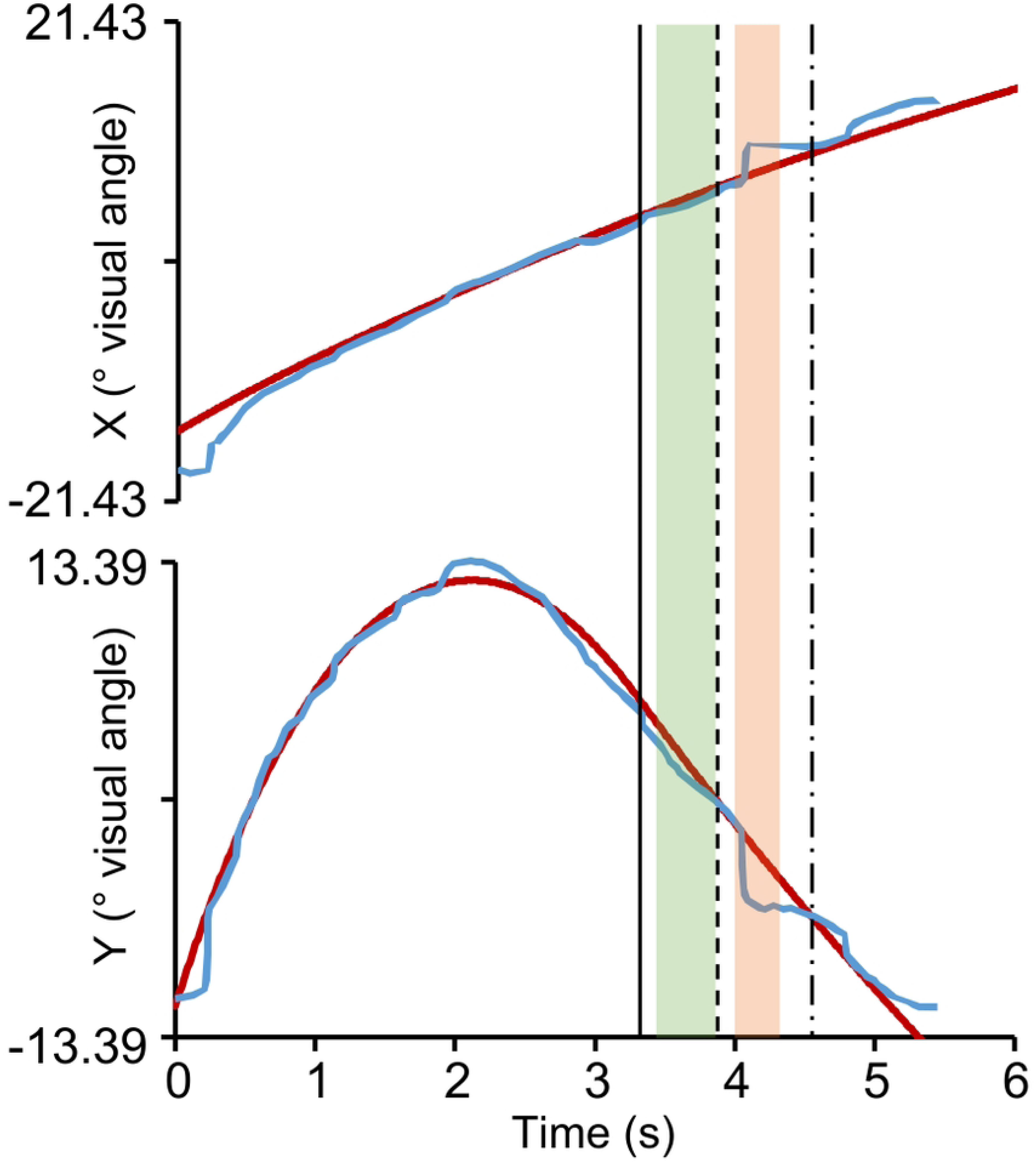
Definition of the *perturbation* and *occlusion* windows for the analysis of the oculomotor behavior. Horizontal and vertical monocular eye-traces (blue) and target trajectory components (red) are illustrated. The vertical solid line represents the perturbation temporal marker and the green semitransparent rectangle the *perturbation* window. Similarly, the dashed vertical line represents the occlusion onset and the orange semitransparent rectangle the *occlusion* window. Finally, the dash-dotted vertical line represents the reappearance of the target.

Since ocular tracking included smooth pursuit bouts alternated with saccadic movements, we derived from the eye traces several indexes related to the saccadic and the smooth pursuit movements. Saccadic movements were detected by using combined eye-velocity (> 30°/s) and eye-acceleration (> 2,000°/s^2^) thresholds (37,39,61). However, saccadic movements that produced post saccadic eye-target distances larger than 4° of visual angle were discarded, as they diverted much the gaze from the target trajectory and, likely, they were not related to the ocular tracking task (37).

From the valid saccades during the perturbation and the occlusion windows we extracted two indexes: the post saccadic error and the saccadic frequency. The post saccadic error was defined as the distance in visual degrees between the eye position after a saccade and the target, averaged among the first 8 ms (4 samples) after the end of a saccade (37). The saccadic frequency was defined as the number of saccades per second.

Smooth pursuit bouts were identified from eye traces after saccades removal, as time epochs with mean eye-to-target distance within 4° of visual angle and mean ratio between eye and visual target velocities comprised between 0.25 and 1.80 (37). Moreover, we considered only smooth pursuit bouts lasting at least 250 ms in the *perturbation* window and at least 100 ms in the *occlusion* window.

From the smooth pursuit bouts we derived the time-lag τ and the gain of the pursuit as defined by Mrotek and Soetching (2007) (48. See also 37,39). Briefly, τ is the lag/lead in milliseconds of the eye relative to the visual target, with positive and negative values indicating that eye movements lagged or were ahead of the target, respectively. The smooth pursuit gain measures the projection of the eye velocity vector on the target velocity vector and, thus, it is a non-dimensional variable. By this definition, a gain of 1 occurs if the eyes move at the same speed and in the same direction as the target, whereas if the eyes move at the target speed but perpendicularly, or if they are stationary (i.e., the subject is fixating) the gain is equal to 0. τ and gain values were computed for each 2 ms data samples and then averaged over the pursuit bout.

### Manual interception analysis

To characterize the interceptive behavior in the Oculo-manual task, we defined three indexes related to the button press responses and to the mouse displacements: the timing error, the positional error and the velocity of the mouse cursor at the time of the button press (velocity at response). The timing error was defined as the temporal difference in milliseconds between the time of the button press and the nominal interception time (i.e. the time at which the target reached the same height in the visual scene as the center of the white circle, which delimited the valid interception area). Negative and positive values indicated anticipated (early) and delayed (late) interceptive responses, respectively. The positional error represented the distance in visual angle between the horizontal location of the white circle at the time of the interceptive response and the horizontal position at which the target trajectory reached the same height as the center of the white circle. Negative values indicated underestimation of the target landing locations whereas positive values indicated their overestimation. The velocity at response was measured in visual angle sec^-1^ with positive and negative values indicating, respectively, rightward and leftward ongoing displacement of the outfielder at the time of the button press response.

### Statistical analyses

Given that the main focus of the study was assessing the effects of the naturalness of the visual scene and of the concurrent execution of the interceptive task on the ocular tracking, we averaged the index values across trials with different initial velocities, perturbation durations and occlusion durations, that is, the manipulations introduced in the experimental design merely to increase the number of different experimental conditions (see above the description of the target trajectories). Thus, for each oculomotor index we pooled, across experimental subjects, the mean values for the reduced number of experimental conditions resulting from collapsing data with respect to target initial velocity, perturbation and occlusion interval, and submitted the resulting datasets to full factorial mixed ANOVA models with the following predictors: visual Scenario (2 levels: Pictorial | Neutral), Gravity Level (3 levels: 0g | 1g | 2g) as *within-subject* factors; and Scenario Order (2 levels: Pictorial first | Neutral first), as well as behavioral Task (2 levels: Oculomotor | Oculo-manual) as *between-subjects* factors. The significance cut-off for main and interaction effects was set to p < 0.01, Greenhouse-Geisser corrected. Moreover, to facilitate interpretation of the significant ANOVA factors, we performed pairwise comparisons between factor levels by using either paired t-tests or two-sample t-tests for dependent or independent measures, respectively (significance cut-off p < 0.01, Bonferroni corrected for multiple comparisons). Consistent with current recommendations (62), we applied restrictive significance cut-offs for the ANOVA effects and the pairwise comparisons to reduce the potential impact of Type I errors, considering also the presence of between subject factors and their interactions in the ANOVA models that could be prone to effects of intergroup variability.

Alike the oculomotor indexes analyses, also for the analyses describing the interceptive performance (Timing error, Positional error, Velocity at response) we used the same reduced set of experimental conditions. The datasets resulting from pooling the mean values of the manual interception indexes across participants to the Oculo-manual task were submitted to full factorial mixed ANOVA models with visual Scenario (2 levels: Pictorial | Neutral) and Gravity Level (3 levels: 0g | 1g | 2g) as *within-subject* factors, and Scenario Order (Pictorial first | Neutral first) as *between-subjects* factor. The significance cut-off for main and interaction effects was set to p < 0.01, Greenhouse-Geisser corrected. We compared differences between levels of the significant factors by means of paired t-tests or two-sample t-tests for dependent or independent measures respectively (significance cut-off p < 0.01, Bonferroni corrected for multiple comparisons).

## RESULTS

We studied the effects of manipulating the degree of realism of the visual scene and of the concurrent execution of a manual interception task on the ocular tracking performance of two groups of participants, one group performing only ocular tracking of the moving target, the other group performing also a manual interception task. Eye tracking performance in the two subject groups was assessed for two distinct temporal windows: the perturbation window, related to the manipulation of the target gravity acceleration; the occlusion window during which vision of the target motion was prevented.

### Ocular tracking performance during the perturbation window

In general, ocular tracking indexes during perturbation window were strongly influenced by factors related to the realism of the visual scene (i.e. the gravity level and the type of visual scenario) and to the type of task performed by participants (see Figure 3 and Table 1).

**Figure 3.**
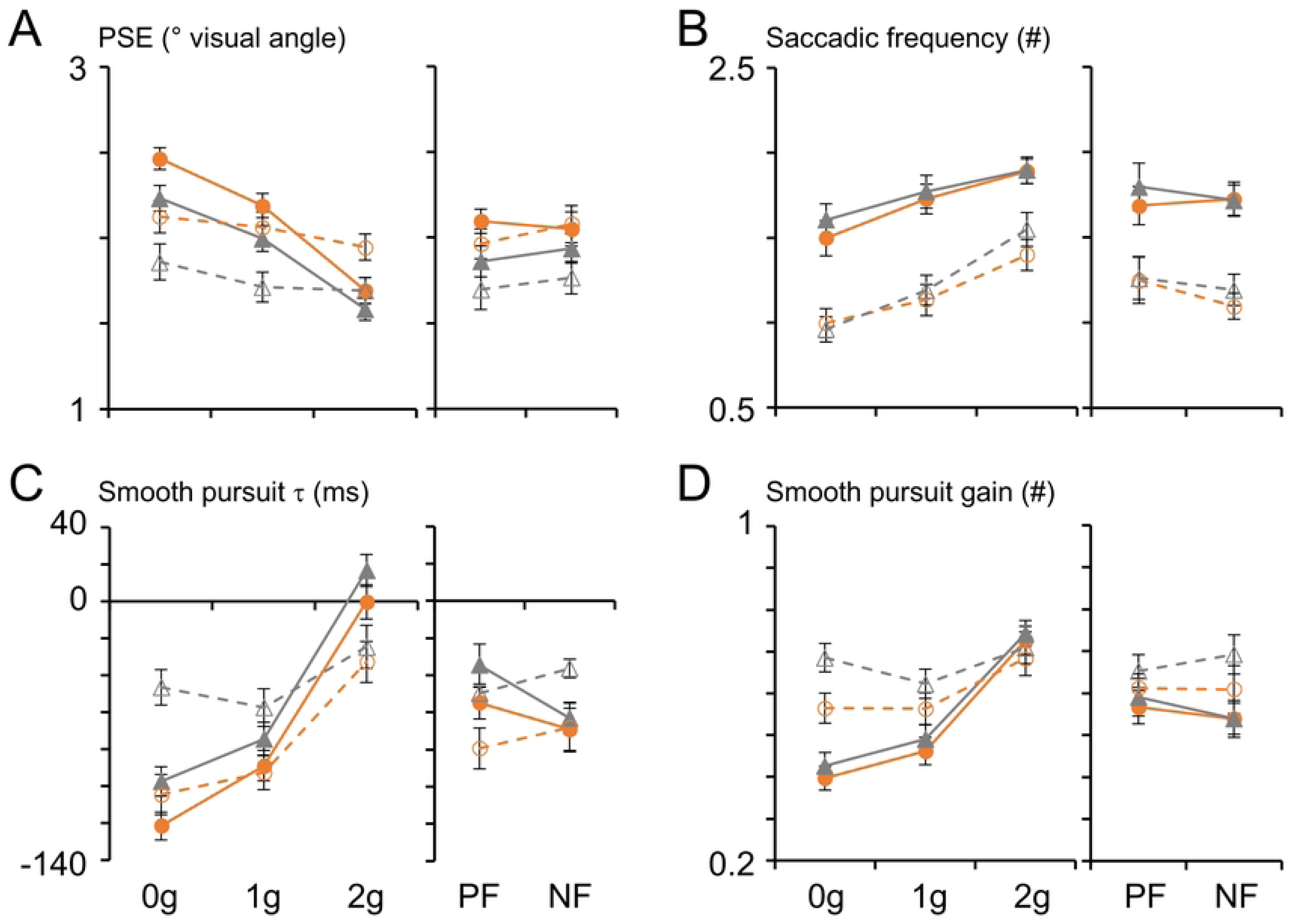
Saccadic and smooth pursuit performance during the *perturbation* window. Each panel consists of two subplots. The first subplot shows the mean (± SEM) oculomotor parameters measured during ocular tracking of the visual targets with either the Pictorial Scenario (orange and circle markers) or the Neutral Scenario (dark grey and triangular markers). Data collected from subjects performing the Oculomotor Task are denoted by filled markers and solid lines, whereas those collected from subjects performing the Oculo-manual Task are denoted by open markers and dashed lines and are grouped across the three law of motion values (0g, 1g and 2g). The second subplot groups the same data according to the order of presentation of the two scenarios averaged across gravity levels (Pictorial first and Neutral first), maintaining the same color coding. A) PSE, post saccadic error. B) Saccadic frequency. C) Smooth pursuit τ. D) Smooth pursuit gain. PF, Pictorial first; NF, Neutral first.

**Table 1.** Results of repeated measures ANOVA on the parameters characterizing saccadic eye movements and SPEMs during the *perturbation* window (in bold p < 0.010, Greenhouse-Geisser corrected). Although the p-values reported are corrected, to simplify the layout of the table and improve its readability, degrees of freedom (dof) are reported uncorrected, once for all the ANOVA tests. S, Scenario; GL, Gravity Level; SO, Scenario Order; T, Task; PSE, post saccadic error; SF, saccadic frequency.

| Factor | dof | PSE | | SF | | $\tau$ | | gain | |
| --- | --- | --- | --- | --- | --- | --- | --- | --- | --- |
|  |  | F | p | F | p | F | p | F | p |
| S | 1,65 | 39.9 | **< 0.001** | 2.0 | 0.158 | 41.4 | **< 0.001** | 16.1 | **< 0.001** |
| GL | 2,130 | 60.6 | **< 0.001** | 42.4 | **< 0.001** | 124.2 | **< 0.001** | 86.8 | **< 0.001** |
| SO | 1,65 | 0.2 | 0.630 | 0.4 | 0.547 | 0.1 | 0.794 | < 0.1 | 0.882 |
| T | 1,65 | 1.9 | 0.174 | 25.5 | **< 0.001** | < 0.1 | 0.929 | 5.1 | 0.027 |
| S*GL | 2,130 | 1.8 | 0.181 | 0.3 | 0.744 | 10.2 | **< 0.001** | 6.9 | **0.003** |
| S*SO | 1,65 | 0.4 | 0.509 | < 0.1 | 0.861 | 0.8 | 0.376 | 0.1 | 0.777 |
| S*T | 1,65 | 2.2 | 0.146 | < 0.1 | 0.963 | 3.4 | 0.070 | 3.4 | 0.071 |
| GL*SO | 2,130 | 0.3 | 0.712 | 0.4 | 0.661 | 0.4 | 0.615 | < 0.1 | 0.986 |
| GL*T | 2,130 | 23.5 | **< 0.001** | 2.1 | 0.135 | 20.5 | **< 0.001** | 29.2 | **< 0.001** |
| SO*T | 1,65 | 0.1 | 0.807 | 0.2 | 0.672 | 2.9 | 0.093 | 0.4 | 0.508 |
| S*GL*SO | 2,130 | 0.8 | 0.427 | 6.6 | **0.003** | 0.9 | 0.398 | 1.0 | 0.355 |
| S*GL*T | 2,130 | 1.0 | 0.365 | 3.5 | 0.039 | 5.8 | **0.009** | 4.2 | 0.024 |
| S*SO*T | 1,65 | 2.7 | 0.103 | 1.7 | 0.191 | 1.3 | 0.252 | 2.5 | 0.116 |
| GL*SO*T | 2,130 | 1.9 | 0.165 | 3.0 | 0.061 | 0.3 | 0.675 | 1.4 | 0.250 |
| S*GL*SO*T | 2,130 | 0.3 | 0.666 | 1.9 | 0.154 | 0.5 | 0.533 | 0.8 | 0.444 |

Manipulations of the target acceleration affected significantly both saccadic and smooth pursuit movements, with all ocular tracking indexes showing monotonic relationships with respect to the target acceleration, which accounted for the highly significant main effects of the gravity level (Table 1). In particular, post saccadic errors were highest and smooth pursuit time leads were largest during 0g trials than 1g and 2g trials, suggesting expectation of gravity effects on the target motion (see Figure 3). Indeed, the paired t-tests evaluating the differences between acceleration levels were all statistically significant, both for the post saccadic errors and the τ values (post saccadic error: 0g vs. 1g, t(_68_) = 4.706, p < 0.001; 1g vs. 2g, t(_68_) = 6.808, p < 0.001; 0g vs. 2g, t(_68_) = 8.179, p < 0.001. τ values: 0g vs. 1g, t(_68_) = 3.213, p = 0.006; 1g vs. 2g, t(_68_) = 11.774, p < 0.001; 0g vs. 2g, t(_68_) = 9.795, p < 0.001). Instead, the monotonic trends observed for the saccadic rates and smooth pursuit gains increased with the target acceleration (Figure 3). However, for smooth pursuit gains, significant changes were evident only for 2g compared to 0g (paired t-test: t(_68_) = 10.476, p < 0.001) and 1g trajectories (paired t-test: t(_68_) = 7.525, p < 0.001), which, in turn, evoked similar pursuit gains (paired t-test 0g vs. 1g: t(_68_) = 1.364, p = 0.531).

In addition to the target acceleration, the presence or absence of quasi-realistic pictorial elements in the visual scene strongly influenced the oculomotor behavior. Specifically, significantly higher post saccadic errors, larger ocular pursuit anticipation and lower pursuit gains were evident in the pictorial compared to the neutral scenario. The saccadic frequency, instead, was not affected (see Figure 3 and the statistics of the main effects of Scenario reported in Table 1).

Smooth pursuit indexes showed also significant interaction effects of Gravity Level*Scenario, which accounted for the observation that τ and gain values differences among gravity levels were reduced in the absence of quasi-realistic pictorial cues. Although this two-way interaction did not reach statistical significance level for the Saccadic Frequencies, we found a similar effect, involving also the order with which the two visual scenes were presented (three-way interaction: Scenario*Gravity Level*Scenario Order, See Figure 3B and Table 1). This interaction effect was mostly accounted by the fact that the saccadic frequency increased monotonically with the gravity level only during the first experimental block, which involved the Pictorial Scenario in one group of subjects and the Neutral Scenario in the other. All pairwise tests evaluating saccadic frequency differences between gravity levels in the first session were, indeed, either statistically significant or borderline significant (paired t-tests: Pictorial first, 0g vs. 1g, t(_33_) = 3.369, p = 0.046; 1g vs. 2g, t(_33_) = 3.602, p = 0.025. Neutral first, 0g vs. 1g, t(_34_) = 3.402, p = 0.041; 1g vs. 2g, t(_34_) = 4.417, p = 0.002).

Regarding the influence of the behavioral task on the ocular tracking behavior, a statistically significant main effect emerged only for the saccadic frequency, which was higher in subjects performing the Oculomotor task. We found, however, statistically significant two-way interaction effects of Gravity Level*Task for all ocular tracking indexes, which accounted for the observation that differences among gravity levels were generally more pronounced in the subject group performing the Oculomotor Task (see Table 1).

For the τ values, significant fraction of the variance was explained also by the three-way interaction Scenario*Gravity Level*Task. This interaction effect was determined by the fact that the slopes of the monotonic trends of the τ values with respect to the gravity levels were different between the two subject groups performing either the Oculo-manual or the Oculomotor task only with the Neutral Scenario. In fact, during the block with the Neutral Scenario, τ values were comparable between the two groups in response to 1g trials (two-sample t-test: t(_67_) = 1.251, p = 1.000) and 2g trials (two-sample t-test: t(_67_) = 2.886, p = 0.126), whereas significant differences between the two subject groups were evident with 0g trials (two-sample t-test: t(_67_) = 4.129, p = 0.003). Instead, mean τ values between subject groups were not significantly different with the Pictorial Scenario.

### Ocular tracking during the occlusion window

During the occlusion window, ocular tracking was primarily influenced by factors related to the degree of realism of the visual stimuli, although with some differences compared to the *perturbation* window (see Figure 4 and Table 2). The gravity level and the type of visual scene were the strongest factors affecting the oculomotor indexes, as signified by their highly significant main effects reported for all oculomotor indexes in Table 2. For the post saccadic error, τ and gain values the main effects of gravity level were accounted for by monotonic trends with respect to the visual target acceleration, akin to those described above for the *perturbation* window (Figure 4). Unlike the *perturbation* window, the mean effect of saccadic frequency was not related a monotonic trend with respect to the target acceleration, because saccadic frequency values were lower during 1g compared to 0g trials (paired t-test: t(_68_) = 3.683, p = 0.001) but not significantly different between 1g and 2g trials (paired t-test: t(_68_) = 1.885, p = 0.191).

**Figure 4.**
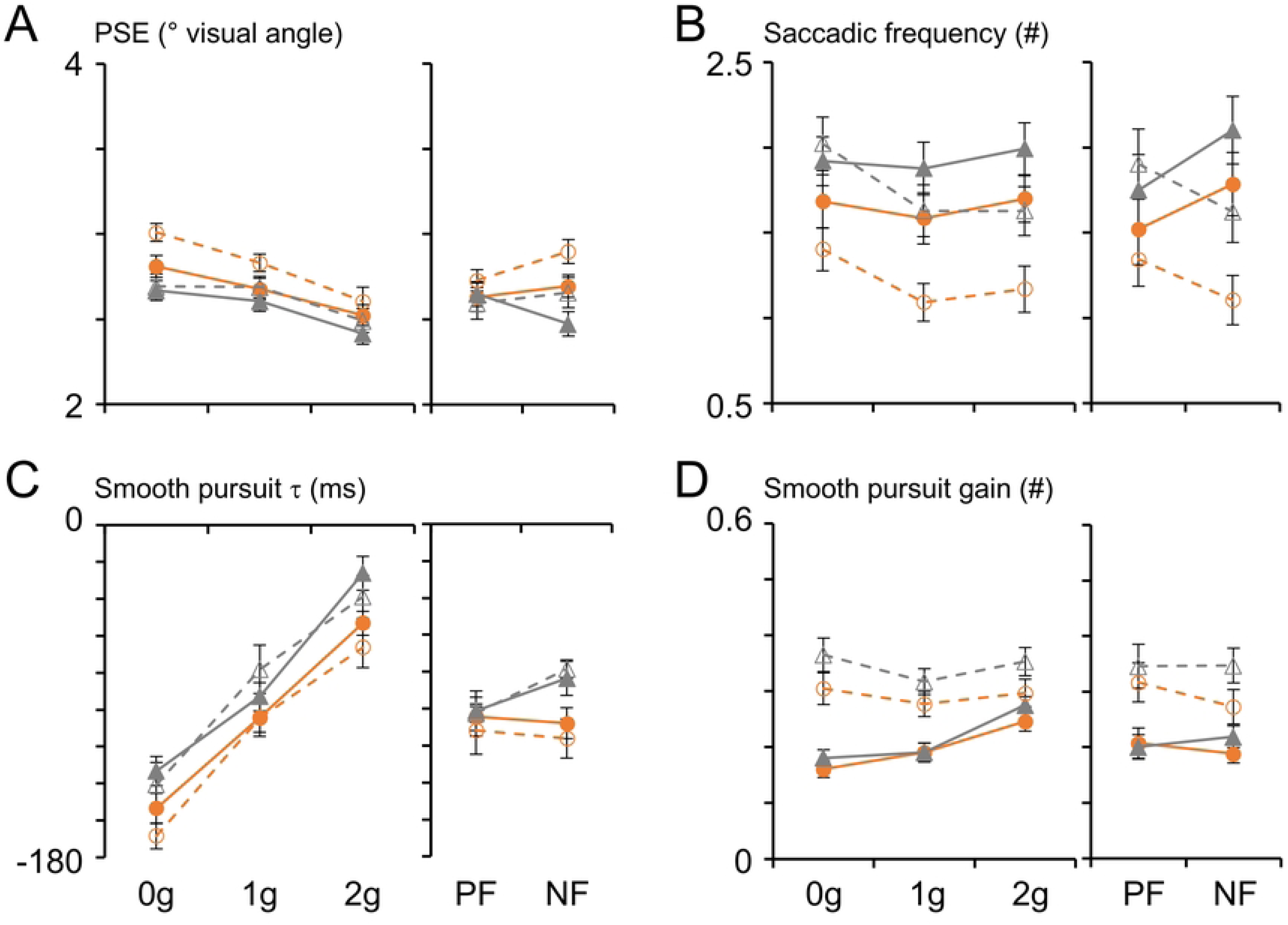
Ocular tracking performance during the *occlusion* window. Same layout as Figure 3.

**Figure 5.**
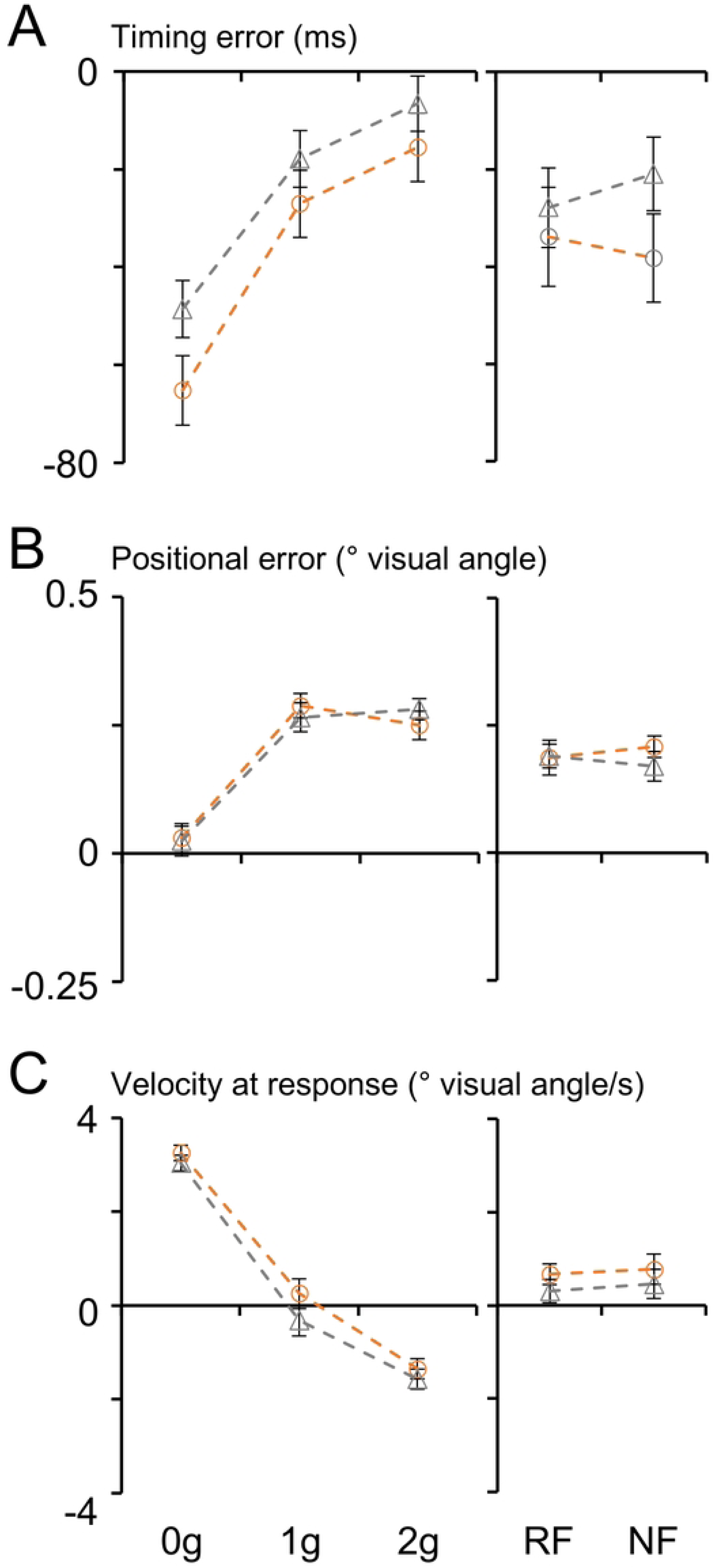
Interception performance in the Oculo-manual Task. The orange dashed lines and the open circles denote the interception parameters’ values (mean ± SEM) obtained with the Pictorial Scenario; the grey dashed lines and the open triangles those obtained with the Neutral Scenario. A) Timing error. B) Positional error. C) Velocity at Response. Same layout as Figure 3 and 4.

**Table 2.** Results of repeated measures ANOVA on the parameters characterizing saccadic eye movements and SPEMs during the *occlusion* window (in bold p < 0.010, Greenhouse-Geisser corrected). Although the p-values reported are corrected, to simplify the layout of the table and improve its readability, degrees of freedom (dof) are reported uncorrected, once for all the ANOVA tests. S, Scenario; GL, Gravity Level; SO, Scenario Order; T, Task; PSE; post saccadic error; SF, saccadic frequency.

| Factor | dof | PSE | | SF | | $\tau$ | | gain | |
| --- | --- | --- | --- | --- | --- | --- | --- | --- | --- |
|  |  | F | p | F | p | F | p | F | p |
| S | 1,65 | 22.8 | **< 0.001** | 53.3 | **< 0.001** | 37.9 | **< 0.001** | 30.2 | **< 0.001** |
| GL | 2,130 | 39.2 | **< 0.001** | 7.0 | **0.005** | 245.7 | **< 0.001** | 20.6 | **< 0.001** |
| SO | 1,65 | 0.1 | 0.719 | < 0.1 | 0.902 | 0.9 | 0.350 | 0.2 | 0.671 |
| T | 1,65 | 2.2 | 0.143 | 2.7 | 0.107 | 0.3 | 0.613 | 17.1 | **< 0.001** |
| S*GL | 2,130 | 3.9 | 0.024 | 0.4 | 0.654 | 0.9 | 0.409 | 3.1 | 0.052 |
| S*SO | 1,65 | 6.8 | 0.011 | 0.1 | 0.817 | 6.2 | 0.016 | 9.7 | **0.003** |
| S*T | 1,65 | 2.0 | 0.163 | 5.7 | 0.020 | 1.1 | 0.297 | 8.3 | **0.005** |
| GL*SO | 2,130 | 1.1 | 0.336 | 2.9 | 0.076 | 0.3 | 0.735 | 0.1 | 0.855 |
| GL*T | 2,130 | 0.2 | 0.805 | 5.5 | 0.012 | 2.9 | 0.071 | 19.1 | **< 0.001** |
| SO*T | 1,65 | 1.0 | 0.312 | 2.5 | 0.121 | < 0.1 | 0.865 | 0.2 | 0.692 |
| S*GL*SO | 2,130 | 1.5 | 0.218 | 8.6 | **< 0.001** | 1.2 | 0.292 | 2.1 | 0.131 |
| S*GL*T | 2,130 | 1.5 | 0.237 | 1.5 | 0.219 | 0.7 | 0.493 | 0.3 | 0.709 |
| S*SO*T | 1,65 | 2.1 | 0.155 | 0.3 | 0.584 | 1.1 | 0.303 | 0.8 | 0.376 |
| GL*SO*T | 2,130 | 1.0 | 0.356 | 0.1 | 0.848 | 0.7 | 0.465 | 1.1 | 0.323 |
| S*GL*SO*T | 2,130 | 5.6 | **0.005** | 1.0 | 0.371 | 4.1 | 0.021 | 0.2 | 0.842 |

Regarding the statistically significant main effects of the type of visual scene on the oculomotor parameters, post saccadic errors, as well as smooth pursuit τ and gain values, showed the same trends observed during the *perturbation* window, namely, longer time leads, lower pursuit gains and higher post saccadic errors in the Pictorial compared to the Neutral Scenario (Figure 4). Unlike the *perturbation* window, however, the saccadic frequency was lower with the Pictorial compared to the Neutral Scenario. The effects of visual scene and target acceleration on the saccadic frequency were influenced further by the order with which the Pictorial and the Neutral Scenario were presented (see the statistically significant three-way interaction Scenario*Gravity Level*Scenario Order in Table 2, also see Figure 4B). This interaction effect was mostly accounted for by the lower saccadic frequency in the Pictorial Scenario compared to the Scenario observed in 0g (paired t-test: t(_33_) = 4.748, p < 0.001) and 1g trials (paired t-test: t(_33_) = 4.375, p = 0.003) in subjects who experienced first the Pictorial Scenario, and in 1g (paired t-test: t(_34_) = 4.743, p < 0.001) and 2g trials (paired t-test: t(_34_) = 6.514, p < 0.001) in subjects who experienced first the Neutral Scenario.

Analogously, for the smooth pursuit gain, we found a significant two-way interaction Scenario*Scenario Order, which accounted for the smaller pursuit gains observed with the Pictorial compared to the Neutral Scenario (paired t-test: t(_34_) = 6.548, p < 0.001) only in participants who experienced the Pictorial Scenario in the second block. In fact, participants experiencing the Pictorial Scenario in the first block did not show significant pursuit gain changes between the two scenarios.

Unlike the perturbation window, where we reported main and interaction effects of the type of behavioral task for several oculomotor indexes, during the occlusion window, the influence of the behavioral task was evident primarily on the pursuit gain and, more marginally, on the post saccadic error. In fact, the only statistically significant main effect of task was observed with the pursuit gain, which was higher, on average, in subjects performing the Oculo-manual compared to those performing the Oculomotor Task (Table 2, see also Figure 4D). The effect of the behavioral task on the pursuit gain was influenced further by the type of visual scene and by the target acceleration level, as indicated by the statistically significant two-way interactions Scenario*Task and Gravity Level*Task reported in Table 2. The Scenario*Task interaction was related to the observation that while the pursuit gains of subjects performing the Oculo-manual Task were significantly lower with the Pictorial than with the Neutral Scenario (paired t-test: t(_31_) = 5.835, p < 0.001), they were not different between scenarios in subjects performing the Oculomotor Task (paired t-test: t(_36_) = 1.399, p = 0.682). The Gravity Level*Task interaction accounted for the different distributions of pursuit gain values with respect to the target acceleration in the two groups of subjects performing either the Oculo-manual or the Oculomotor task. In fact, clear monotonic trends of the pursuit gain with the gravity lever were observed only in subjects performing the Oculomotor Task (paired t-test: 0g vs. 1g, t(_36_) = 1.896, p = 0.594; 1g vs. 2g, t(_36_) = 7.512, p < 0.001; 0g vs. 2g, t(_36_) = 8.011, p < 0.001) but not in those performing the Oculo-manual Task (paired t-test: 0g vs. 1g, t(_31_) = 3.188, p = 0.029; 1g vs. 2g, t(_31_) = 3.303, p = 0.022; 0g vs. 2g, t(_31_) = 0.538, p = 1.000; see also Figure 4D).

Finally, a small influence of the type of behavioral task was evident also for the post saccadic error with a statistically significant four-way interaction effect of Scenario*Gravity Level*Scenario Order*Task (Figure 4A and Table 2).

### Interception performance

We characterized subjects’ interceptive behavior by way of mixed ANOVAs on the Timing error, Positional error and Velocity at response (see Figure 6 and Table 3). A common finding emerging from these analyses was that Gravity Level represented the major factor accounting for the interceptive performance across experimental conditions, as indicated by the highly significant main effects reported for all three indexes in Table 3. In general, the changes in the interceptive performance related to the gravity level, seemed compatible with the use of a priori knowledge of gravity effects on the target motion. For example, the distribution of Timing error values across gravity levels denoted significantly stronger anticipation of the interceptive responses to 0g targets compared to accelerated 1g (paired t-test: t(_31_) = 13.854, p < 0.001) and 2g (paired t-test: t(_31_) = 17.149, p < 0.001) targets, compatible with the a priori expectation of gravity. In addition to the Gravity Level, Timing error values were significantly influenced by the type of Scenario, as participants produced earlier interceptive responses with the Pictorial Scenario compared to the Neutral Scenario, and by the three-way interaction Scenario*Gravity Level*Scenario Order (Table 3).

**Table 3.** Results of repeated measures ANOVA on the parameters characterizing the interceptive performance (in bold p < 0.010, Greenhouse-Geisser corrected). Although the p-values reported are corrected, to simplify the layout of the table and improve its readability, degrees of freedom (dof) are reported uncorrected, once for all the ANOVA tests. TE, Timing error; PE, Positional error; VaR, Velocity at response

| Factor | dof | TE |  | PE |  | VaR |  |
| --- | --- | --- | --- | --- | --- | --- | --- |
|  |  | F | p | F | p | F | p |
| S | 1,30 | 11.0 | **0.002** | < 0.1 | 0.959 | 16.8 | **< 0.001** |
| GL | 2,60 | 219.2 | **< 0.001** | 64.0 | **< 0.001** | 184.7 | **< 0.001** |
| SO | 1,30 | < 0.1 | 0.917 | 0.3 | 0.610 | 0.1 | 0.737 |
| S*GL | 2,60 | 3.3 | 0.061 | 1.8 | 0.184 | 6.5 | **0.005** |
| S*SO | 1,30 | 2.8 | 0.106 | 1.1 | 0.296 | 0.1 | 0.781 |
| GL*SO | 2,60 | 3.0 | 0.064 | 0.2 | 0.851 | 0.7 | 0.442 |
| S*GL*SO | 2,30 | 9.3 | **0.001** | 25.3 | **< 0.001** | 1.9 | 0.167 |

Positional differences across experimental conditions were much smaller than those observed for the Timing error (see Figure 6B). Nevertheless, we found a significant main effect of the Gravity Level, related to the slight overestimation of the responses to accelerated 1g (one sample t-test: t(_31_) = 11.327, p < 0.001) and 2g (one sample t-test: t(_31_) = 13.162, p < 0.001) targets, compared to the responses to 0g targets, which were not different from zero (one sample t-test: t(_31_) = 1.261, p = 0.217). Alike the interceptive timing, Positional error values were influenced significantly also by the three-way interaction Scenario*Gravity Level*Scenario Order.

A monotonic relationship with the Gravity Level emerged clearly also with the values of the Velocity at response, accounting for the highly significant main effect of this factor (Table 3, Figure 6C). Velocity at response values were close to zero, on average, when subjects intercepted 1g targets (one-sample t-test: t(_31_) = 0.108, p = 0.915), denoting that the cursor was firmly in place at the time of the response. Instead, with 0g and 2g targets, Velocity at response values were positive (one-sample t-test: t(_31_) = 19.391, p < 0.001) and negative (one-sample t-test: t(_31_) = 6.968, p < 0.001), respectively, indicating that subjects, at the time of the button press, were making corrections of the mouse cursor position in opposite directions for these two trajectories, compatible with expectation of gravity effects on the target motion.

Velocity at response values were also influenced by the type of Scenario, with slightly higher values when the task was performed with the Pictorial than with the Neutral Scenario (significant main effect of Scenario in Table 3). Finally, the statistically significant two-way interaction Scenario*Gravity Level, indicated that differences in Velocity at response values between the two scenarios were statistically significant during 1g trials (paired t-test: t(_31_) = 4.163, p = 0.002), but not in 0g and 2g trials (paired t-tests: t(_31_) = 2.939, p = 0.056; and t(_31_) = 2.416, p = 0.196, respectively).

## DISCUSSION

In this study, we reported on the influence of the naturalness of a visual scene and of the presence or absence of concurrent manual interception of the visual target on the ocular tracking performance.

### Effects of naturalness of the visual environment on ocular tracking

One type of manipulation of the naturalness of the visual environment was imposing sudden changes of the acceleration profile of the visual targets, thereby making their kinematics less compatible with the effects of natural gravity. The rationale for this manipulation stemmed from earlier evidence that a priori knowledge of gravity effects can be used effectively for the predictive control of manual interceptions as well as of eye movements (for a review, see 25). We hypothesized that if ocular tracking control mechanisms relied on an expectation of gravity effects, the oculomotor indexes characterizing saccadic and smooth pursuit kinematics would show systematic dependence on the acceleration levels imposed to the visual targets, similar to what reported for the interception of objects under comparable experimental conditions (21,23,24,63). Conversely, if control of ocular tracking was based on mechanisms relying primarily on sensory information, eye movement kinematics would reflect merely the changes of the target speed profiles imposed by the experimental manipulations of the target acceleration (64–67). We found that ocular tracking was strongly influenced by the target acceleration, regardless of the availability of visual information about target motion. Ocular tracking indexes showed, in fact, significant monotonic trends with respect to the target acceleration, denoting stronger anticipation of smooth pursuit movements and higher post saccadic errors during 0g trajectories compared to accelerated 1g and 2g trajectories, as well as higher pursuit gains and saccadic rates for accelerated trajectories. Like our previous studies adopting similar experimental conditions, the differences in the behavioral responses between 1g and 2g accelerated motion did not seem very consistent across oculomotor indexes or the 2g even bettered 1g responses (21,37,39). We attributed this issue to the poor sensitivity of the visual system to retinal acceleration (68,69), and to the possibility that visual context information provided by the pictorial scenario could not be sufficient to scale adequately retinal acceleration to external world acceleration. Thus, overall, the present results were consistent with earlier reports suggesting that the changes in the ocular tracking performance observed following manipulations of the visual target acceleration might reflect a priori expectations of gravity effects (37,39,70,71).

In addition to the parametric changes of the target acceleration, we manipulated the realism of the visual background and found that visual context information influenced greatly ocular tracking, regardless of the availability of visual motion information. The changes in the oculomotor performance were consistent with those reported by a previous study from our laboratory employing a similar set of experimental conditions (37). In the current study, post saccadic errors were larger and τ values more negative in the pictorial compared to the neutral scenario during both the perturbation and the occlusion windows. These results may be compatible with a greater level of anticipation –and, therefore, a higher degree of predictability– of the target’s motion when visual cues aid in interpreting its trajectory (72). Noteworthy, this trend was evident also when the target motion was visually occluded, as information gathered while the target was visible, along with a priori knowledge, may have been used to maintain an anticipatory strategy in the absence of visual feedback about the target motion (73,74).

Smooth pursuit gain values were lower in the pictorial than in the neutral scenario during both perturbation and occlusion windows. This result appears in line with previous findings that textured backgrounds can fragment eye tracking, by reducing the pursuit gain and increasing the number of saccades (75,76). The increased saccadic rate found by Delle Monache and colleagues (2019), however, was not replicated by the present study, since during the occlusion window we observed higher saccadic rates with the neutral scenario, consistently across subject groups performing either the Oculo-manual or the Oculomotor Task. Moreover, in this earlier study we reported an interaction effect, during the perturbation window, between the gravity level and the realism of the visual scene that was not replicated by the present study, where we observed a certain influence of gravity level in both scenarios, although to a slightly different extent.

We may explain these differences between the earlier and the current study by assuming that participants were attempting to predict the target reappearance beyond the occluder, rather than tracking it, since the contour of the occluder cued where the target would reappear. In fact, in our previous study –where targets were not concealed behind a visual occluder but simply vanished for the same variable intervals as the present study– we found higher pursuit gains.

In other words, the occluder could have represented an attentional *attractor*, by providing cues about the onset and the spatial extent of the occlusion. By using these cues, participants may have shifted their gaze earlier to the occluder, rather than approaching it at the same speed as the target. Compatible with this view, Villavicencio and colleagues (2024) (77) showed that knowing the location of the occlusion onset during an interceptive task could increase the number of saccades towards the occluder, whereas if the occlusion onset was unpredictable, this behavior did not occur. Moreover, they observed higher pursuit gains when the occlusion onset was unpredictable compared to when it was predictable, likely because of a strong reliance on sensory information than on predictive mechanisms. Indeed, the findings reported by Villavicencio and colleagues (2024) (77) may help reconciling the differences we found between our previous study and the current one, since in the current study the onset of the occlusion was much more predictable than in the previous study. Thus, the predictability of the occlusion in the current experiment might have biased the saccadic behavior reducing consistently the potential effects of the manipulations of the visual scenario that emerged in the earlier study. In addition, it might explain also the higher gains we observed during the perturbation window in the previous compared to the present study.

Another possibility to consider is that the occluder may have also worked as a *distractor*, independently of the type of visual scenario. It is known, in fact, that distractors can disrupt attention and influence saccadic movements (78–80), thereby potentially inducing saccadic errors. Smooth pursuit indexes (τ and gain values), however, were consistent with our previous study, showing stronger dependence on the target’s law of motion in the pictorial than in the neutral scenario. The discrepancy that saccadic and smooth pursuit eye movements were affected differently by the interaction of the target kinematics with the type of visual scenario could be, in part, explained by the fact that, while distractors can also affect smooth pursuit (81), their influence is more pronounced during the initial open-loop phase. The influence of the distractor is, in fact, suppressed by selective attention mechanisms during the subsequent closed-loop phase driven by sensory signals, especially when, like in our study, the distractor’s position is rather predictable (82).

It must be pointed out, however, that, in the present experiment, the occluder initially fell in the peripheral vision, ruling out any distractor effect earlier on in the trial. As the target reached the perturbation window not less than 1.8 seconds later –i.e. long after the 450 ms open-loop phase subject to the influence of distractors (83)– its distance from the occluder was at least ∼6.5 visual degrees, implying that the occluder image was still projected to the parafoveal region (84). Thus, we may suggest that any potential distractor effects on the smooth pursuit movements were likely suppressed by the time the target reached the perturbation window, perhaps explaining why smooth pursuit indexes were not as affected by the introduction of the graphic occluder.

### Effect of the naturalness of the visual environment on the interceptive response

With respect to the interceptive performance of the participants who performed the Oculo-manual Task, we found general agreement with previous findings based on similar manipulations of the visual target law of motion (21,85,86). In fact, timing and spatial errors reflected an underestimation of 0g trajectories compared to accelerated (1g and 2g) trials, a result considered compatible with gravitational expectation (21). The distributions of mouse cursor velocities at the time of button presses were compatible with gravitational expectation as well, by denoting underestimation of 0g trajectories landing positions and overestimation of the 2g ones. Similar to Bosco et al. (2012) (21), we did observe values that were close to zero (i.e. no final corrections on the estimates of the targets landing position) for 1g trials, with evident rightward or leftward corrective movements during 0g and 2g trials, respectively.

### Influence of the concurrent interception task on ocular tracking

The execution of concurrent tasks may influence various aspects of oculomotor control, as clearly shown by the seminal findings by Yarbus (40). Indeed, oculomotor control processes can be affected by the expected outcome of a specific behavioral strategy (87–89), by the nature of the concurrent task (90) and by its computational demands (91,92). Moreover, there is evidence that performing two demanding tasks simultaneously may reduce the quality of ocular tracking (41,43–45). Conversely, oculomotor strategies competing for attentional resources may introduce time gaps in cognitive tasks (93).

With respect to the concurrent execution of an interceptive task, it is known that ocular tracking performance can influence the interception accuracy (52,94), by way of a corollary discharge of the oculomotor commands, exemplifying a direct link between the precision of the ocular tracking and the performance of a specific concurrent task. This is particularly interesting from the perspective of the neural control mechanisms involved. For example, both ocular tracking and manual interceptive control involve predictive mechanisms, but the degree to which predictive information may be shared or weighted differentially between the two control systems is still a matter of debate. In this regard, internal models of the physical properties of the environment appear to contribute critically to the predictive estimates of the timing of interceptive movements in humans (95–97), in non-human mammals (98,99), and even in some invertebrates (100). Similar evidence that predictions about the object motion may be based on internal models of gravity effects has been gathered also for oculomotor control (37–39), suggesting that ocular tracking and manual interception share the same predictive information. However, it is not clear whether this predictive information is engaged with a similar temporal course between the two tasks.

A recent study from our group may give some indication about the time course with which internalized information about gravity may be engaged by the predictive processes underlying the control of interceptive timing (22). In this study, we modelled the interceptive timing performance of subjects intercepting looming stimuli in a quasi-realistic 3D virtual reality setting, with bayesian regression models which included a predictor related to an internal estimate of gravity effects, which could be engaged at successive time-points from the beginning of the visual motion up to 600 ms thereafter. The results of this analysis suggested that subjects relied exclusively on optical information during the first 450 ms of the descending trajectory (i.e., up to 350-650 ms before the nominal interception point, as target motion was comprised between 800 and 1100 ms), after which participants engaged and relied mostly on the gravity prior. In our current study, the changes in oculomotor behavior might be used to infer the time-point at which the transition from sensory information to prior expectations occurred. During the perturbation window, with visual feedback about the target’s position available, the dependence of saccadic and smooth pursuit parameters on expectation of gravity effects was clearly evident in subjects performing only ocular tracking, but it was weaker for participants performing also the target interception (as indicated by Gravity Level*Task p < 0.001 for all oculomotor variables except saccadic frequency, see Figure 3 and Table 1). Later in the trial, during the occlusion window, similar monotonic trends with respect to the gravity level emerged for post saccadic errors and smooth pursuit τ values regardless of the task, even though task-dependent differences persisted for smooth pursuit gain values. These findings imply that participants performing the Oculomotor Task began relying strongly on gravitational expectations at least 950-1200 ms before the target’s nominal landing. In contrast, when interception was also required (as for participants performing the Oculo-manual Task), the transition from sensory information to internal model predictions occurred only after the target occlusion (i.e. not earlier than 400-650 ms before target’s nominal landing). That is, the introduction of the interception task seemed to postpone the engagement of the internal model of gravity until predictive mechanisms were enforced by the visual occlusion of the moving target, as if subjects increased reliance on sensory signals as long as they were available to exploit every bit of incoming information (77,101). Although the obvious differences in the experimental design and analysis make it difficult to compare directly the findings of the two studies, the current analysis of the oculomotor indexes between the two behavioral tasks does suggest a temporal course for the engagement of the internal model of gravity like that implied by earlier study. Thus, oculomotor and manual interception control may rely on a common mechanism that engages predictions of an internal gravitational model with a similar temporal profile when the two tasks are performed concurrently.

## Conclusion

In sum, the current study provided further support to the idea that expectation of gravity effects on the motion of visual objects can influence significantly ocular pursuit, much alike what consistently reported for interceptive movements. By manipulating the type of visual scenes created with computer graphics, we also substantiated earlier findings that pictorial cues informing about the naturalness of the environment can reinforce gravitational expectations, particularly in the absence of visual motion information, which makes more crucial the reliance on built-in predictions of the target’s behavior. Finally, concurrent execution of the ocular tracking and of the interceptive task provided interesting insights on the temporal recruitment of the internalized gravity information, since gravitational expectations were applied consistently –even before predictive mechanisms were enforced by the target occlusion– by subjects performing only ocular tracking, whereas concurrent execution of the two tasks postponed the full recruitment of the internal gravity model to the period of visual occlusion, by increasing the reliance on visual information as long as it was available.

## ACKNOWLEDGEMENTS

The work was supported by the Italian University Ministry (#NEXTGENERATIONEU (NGEU) National Recovery and Resilience Plan (NRRP), project MNESYS PE0000006 – A Multiscale integrated approach to the study of the nervous system in health and disease DN. 1553 11.10.2022), BRIC INAIL (Laborius ID57-2022), U.S. Department of Defense Congressionally Directed Medical Research Program W81XWH1810760 PT170028, by #NEXTGENERATIONEU (NGEU), Ministry of University and Research (MUR), National Recovery and Resilience Plan (NRRP), PRIN 2022B42X54, by the Italian Ministry of Health (RF-2019-12369194 and IRCCS Fondazione Santa Lucia Ricerca Corrente), by #NEXTGENERATIONEU (NGEU), Ministry of Health, National Recovery and Resilience Plan (NRRP), PNRR-MCNT2-2023-12377870, and by the Italian Space Agency, ASI, and the Ministry of University and Research, MUR, under contract n. 2024-5-E.0 - CUP n. I53D24000060005

